# Impact of bacterial microcompartment-dependent ethanolamine and propanediol metabolism on *Listeria monocytogenes* interactions with Caco-2 cells

**DOI:** 10.1101/2021.08.26.457845

**Authors:** Zhe Zeng, Lucas M. Wijnands, Sjef Boeren, Eddy J. Smid, Richard A. Notebaart, Tjakko Abee

**Author notes:** Corresponding author Tjakko Abee.

## Abstract

Bacterial microcompartment (BMC) dependent ethanolamine (*eut*) and propanediol utilization (*pdu*) has recently been shown to stimulate anaerobic growth of *Listeria monocytogenes*. This metabolic repertoire conceivably contributes to the competitive fitness of *L. monocytogenes* in the human gastrointestinal (GI) tract, where these compounds become available following phospholipid degradation and mucus-derived rhamnose metabolism by commensal microbiota. Previous transcriptomics and mutant studies of *eut* and *pdu L. monocytogenes* suggested a possible role of *eut* and *pdu* BMC metabolism in transmission in foods and pathogenicity, but data on a potential role of *L. monocytogenes* interaction with human cells is currently absent. First, we ask which cellular systems are expressed in the activation of *eut* and *pdu* BMC metabolism and the extent to which these systems are conserved between the states. We find common and unique systems related to metabolic shifts, stress and virulence factors. Next, we hypothesize that these common and unique activated cellular systems contribute to a role in the interaction of *L. monocytogenes* interaction with human cells. We present evidence that metabolically primed *L. monocytogenes* with active *eut* and *pdu* BMCs, as confirmed by metabolic analysis, transmission electron microscopy and proteomics, show significantly enhanced translocation efficacy compared to non-induced cells in a trans-well assay using Caco-2 cells, while adhesion and invasion capacity was similar. Taken together, our results provide insights into the possible key cellular players that drive translocation efficacy upon *eut* and *pdu* BMC activation.

## 6.1 Introduction

*Listeria monocytogenes* is a food-borne pathogen responsible for a severe infection called listeriosis, which case-fatality can reach up to 30% in specific high-risk population groups such as the elderly, immunocompromised individuals, fetuses and newborns [1]. Acquisition of this infection is mainly caused by consumption of contaminated food (predominantly ready-to-eat food) [2]. *L. monocytogenes* is found ubiquitously in nature environments such as soil, silage, groundwater, sewage and vegetation [1, 3]. The food-borne pathogen can grow at low temperatures and is very robust to survive in environmental stresses, such as low pH and high salt concentrations [1, 2]. All of these features make the transmission and contamination of *L. monocytogenes* a severe concern for the food industry [2, 4, 5].

Upon ingestion of contaminated food by the host, *L. monocytogenes* needs to overcome acidic conditions in the stomach before reaching the GI tract and subsequent interaction with the mucus-coated epithelial barrier [1, 6]. *L. monocytogenes* has evolved a number of mechanisms to cross various host barriers and survive with its unique intracellular lifestyle [1, 6, 7]. Following the entry into epithelial cells mediated by Internalin A (InlA) and Internalin B (InlB), *L. monocytogenes* is internalized into the vacuole [1, 8]. Subsequently, *L. monocytogenes* produces listeriolysin O (LLO) [6] and two phospholipases, phospholipase A (PlcA) and phospholipase B (PlcB), for vacuolar rupture and escape, which together with Actin (ActA), involved in intracellular motility and cell to cell spread, mediate crucial steps in *L. monocytogenes* pathogenesis [1, 3, 9]. *L. monocytogenes* can survive and duplicate within the cytosol of the host cell and even induce changes in the morphology of host cell organelles, thereby altering their function to promote infection [1, 3]. Notably, alternative routes of entry and translocation have been described including possible roles for so-called Listeria Adhesion Protein (LAP, lmo1634) [10, 11], and listeriolysin/LLO [12]. The bacterium, first used as a model to study cell-mediated immunity, has emerged over the past 20 years as a paradigm in the study of bacterial regulation and host-pathogen interactions and, more recently, interactions with the gut microbiome. (Reviewed in [1, 3]).

Upon arriving in the host gut, *L. monocytogenes* needs to adapt to an environment already rich in commensal microbes and poor in available nutrients. Recent evidences suggest that *Salmonella* and other enteric bacteria conduct cellular organelles called bacterial microcompartments (BMCs) to facilitate their pathogenesis in host gut [13–15]. BMCs are proteinaceous organelles that optimize the utilization of some specific substrates, such as 1,2-propanediol, ethanolamine, rhamnose and choline, with the encased enzymes producing toxic intermediate aldehydes, thereby preventing damage to cells’ DNA and proteins [14, 16]. Substrates for BMC-dependent metabolism like ethanolamine and 1,2-propanediol can be produced in the GI tract by bacterial metabolism of (host) phospholipids and following mucus degradation and rhamnose metabolism, respectively [13, 17, 18]. Enteric pathogens gain a competitive advantage in the intestine by utilizing these substrates, an advantage enhanced by the host inflammatory response (Reviewed in [17]). Previous studies using cell lines and animal models and assessing performance of pathogen wild type and selected mutants in BMC dependent *eut* and *pdu*, pointed to a role in virulence of foodborne pathogens including *Salmonella enterica* Serovar Typhimurium, *Escherichia coli*, and *L. monocytogenes* [17, 19, 20].

The transcriptomics and mutant studies of *pdu* and *eut L. monocytogenes* suggested a possible role in transmission in foods and pathogenicity. Transcriptomics in *L. monocytogenes* showed the upregulated expression of *pdu* and *eut* for growth on vacuum-packed cold smoked salmon[21] and mouse infection model [22]. A deleted *pduD* mutant of the *L. monocytogenes* showed attenuated virulence and faster clearing of *L. monocytogenes* in the GI tract which indicate the importance of *pdu* for the fitness of *L. monocytogenes* during infection in the GI tract [22]. The importance of *eut* and *pdu* activation in stress survival and virulence of *L. monocytogenes* was highlighted in recent reviews [19]. Notably, we recently provided experimental evidence including metabolic and proteomic analysis, combined with transmission electron microscopy, for activation of *L. monocytogenes pdu* and *eut* BMCs and corresponding growth stimulation in anaerobic conditions [23, 24]. Combining our data with previous studies, we hypothesize that priming of *L. monocytogenes pdu* and *eut* BMCs in foods and/or the intestine, may affect interaction with the host epithelial barrier. In this study we quantified adhesion, invasion and translocation efficacy of *pdu*-induced and *eut*-induced *L. monocytogenes* compared to non-induced cells in Caco-2 cell assays. Correlations of virulence results with *L. monocytogenes* metabolic, stress and virulence factors, that show significant differential expression in *pdu*-induced and *eut*-induced cells versus non-induced cells, are discussed.

## 6.2 Materials and Methods

### 6.2.1 Strain and Culture Conditions

*L. monocytogenes* EGDe was used in this study. *L. monocytogenes* EGDe was anaerobically grown at 30°C in Luria Broth (LB) medium supplemented with 50mM 1,2-propanediol or 15mM ethanolamine (EA) as extra carbon source as described before [23, 24]. 20nM Cobalamine B12 was added to induce *pdu* and *eut* cluster as B12 controls riboswitch-based regulator of *pdu* and *eut* cluster [25, 26]. Anaerobic conditions were achieved by Anoxomat Anaerobic Culture System with the environment 10% CO_2_, 5% H_2_, 85% N_2_. LB with 50 mM 1,2-propanediol or 15 mM ethanolamine plus 20 nM vitamin B12 was defined as *pdu*-induced and *eut*-induced condition, respectively, while LB with 50 mM 1,2-propanediol or 15 mM ethanolamine conditions without B12 were defined as *pdu* and *eut* non-induced conditions. *L. monocytogenes* cells were harvested until the late-exponential growth phase (OD600 = 0.4–0.5) for proteomics analysis and Caco-2 infection test.

### 6.2.2 Proteomics

*L. monocytogenes* EGDe cultures were anaerobically grown at 30°C in *pdu/eut* induced and in non-induced conditions. Samples were collected at 12 h of the inoculation and processed as described before [23, 24]. The mass spectrometry proteomics data have been deposited to the ProteomeXchange Consortium via the PRIDE [27] partner repository with the dataset identifier PXD027979 for *eut* induced and in non-induced conditions (retrieved from Zeng Z, et al, 2021 [24]) and identifier PXD028010 for *pdu* induced and in non-induced conditions.

### 6.2.3 Caco-2 adhesion, invasion and translocation

Cultures of Caco-2 cells (human intestinal epithelial cells, ATCC HTB-37), production of differentiated cells in 12-well plates were carried out as described before [25]. *L. monocytogenes* overnight grown cells used for adhesion, invasion and translocation were normalized to obtain cell concentrations of 8.0 ± 0.2 log CFU/ml.

For adhesion and invasion experiments with Caco-2 cells, 1.6 × 105 cells/well were seeded into the 12-well tissue culture plates (Corning Inc. ID 3513). Inoculated 12-well plates were incubated at 37 °C for 12–14 days, with the medium refreshed every 2 days, to establish a confluent monolayer of cells. Adhesion and invasion experiments were started by inoculation with 40 μl of late exponential phase cells of *pdu/eut* induced and non-induced *L. monocytogenes* EGDe resulting in an final inoculum of approximate 6.8 log CFU/well. Then the 12 well plates were centrifuged for 1 min at 175 ×g to create a proximity between the *L. monocytogenes* cells and Caco-2 cells.

For adhesion and invasion enumeration, after 1h anaerobic incubation without gentamicin, *L. monocytogenes* cells that have not adhered to Caco-2 cells, were removed by washing three times with PBS buffer. Half of the wells containing Caco-2 cells were lysed with 1 ml of 1% v/v Triton X-100 in PBS and serially diluted in PBS for quantification of the number of adhered and invaded *L. monocytogenes* cells. The other half of the wells of the Caco-2 cells was subsequently incubated anaerobically for 3 h with 0.3% gentamicin (50 μg/ml, Gibco) to eliminate all extracellular *L. monocytogenes* cells. Thereafter, gentamycin containing medium was removed by washing three times with PBS buffer. The Caco-2 cells were lysed with 1 ml of 1% v/v Triton X-100 in PBS and serially diluted in PBS for quantification of the number of invaded *L. monocytogenes* EGDe cells.

For translocation experiments, 0.8 x 105 Caco-2 cells per mL were seeded ThinCert PET inserts Greiner Bio-One 665640) for 12–14 day differentiation at 37 °C. Prior to the translocation assay wells and inserts were washed three times with PBS, and placed in TCM without gentamycin and fetal bovine serum. Translocation was started by adding 20 μl of late exponential cells of *pdu/eut* induced and non-induced *L. monocytogenes* EGDe into the inserts, resulting in an inoculum of approximately 6.5 log CFU/well. After centrifuging the Transwell plates for 1 min at 175x g, the plates were incubated 2 hrs anaerobically. After incubation, the inserts were removed with a sterile forceps and discarded. The contents in the wells was collected for quantification of the translocated number of *L. monocytogenes* EGDe cells.

### 6.2.4 BMC gene loci analysis

The Hidden Markov Models (HMMs) of two BMC shell protein domains listed as Pf00936 and Pf03319 were retrieved from the Pfam database to predict BMC shell proteins in *L. monocytogenes* EGDe as described before [23, 24]. Shell proteins were predicted by a HMM search using the HMMER package and a local protein database of *L. monocytogenes* EGDe genome [28]. All hits with an e-value less than or equal to 1e-05 that correspond to a genomic record from Genbank, RefSeq, EMBL, or DDBJ databases were accepted as BMC shell protein homologs. The annotation from InterPro are taken to fulfill the annotation of some unannotated proteins in BMC loci. Details in Supplementary Table1.

### 6.2.5 Venn analysis and STRING networks analysis

The protein IDs of significantly changed proteins from Supplementary Table 2 and 3 were uploaded to the BioVenn online server [29] taking the default setting to generate Venn diagrams. Overlapping proteins from the Venn diagram were transferred to the STRING online server[30] for multiple proteins analysis of functional interaction using sources such as co-expression, genomic neighbourhood and gene fusion.

### 6.2.6 Pathway visualization of protein expression via Pathview

The UniProt protein IDs from Supplementary Table 2 and Supplementary Table 3 were collected and retrieved to Entre IDs. A list of Entrez IDs, protein expression indicated by LFQ intensity (Supplementary Table 4 and Supplementary Table 5) was mapped to the *L. monocytogenes* EGDe KEGG pathway database using the tool Pathview (R version 3.2.1) [31]. The box represent genes and the different color indicates level of expression with default setting.

Statistical analyses were performed in Prism 8.0.1 for Windows (GraphPad Software). As indicated in the figure legend, Statistical significances are shown in ***, P<0.001; *, P<0.05; ns, P>0.05 with Holm-Sidak T-test.

## 6.3 Results

### 6.3.1 *pdu* and *eut* BMC gene loci organization and core metabolic reactions

The genes encoding conserved BMC shell proteins and core enzymes are located in respective *pdu* and *eut* clusters, and corresponding metabolic reactions in *L. monocytogenes* EGDe are shown in Figure 1. Briefly, the *pdu* cluster contains seven BMC shell proteins PduTUABKJN and the *eut* cluster contains five BMC shell proteins EutLKMN and Lmo1185 (Figure 1A and Supplementary Table 1) as previously described [23, 24]. BMC substrates 1,2-propanediol and ethanolamine, enter respective BMCs and are degraded via their signature enzymes, propanediol dehydratase PduCDE and ethanolamine ammonia lyase EutBC, into corresponding aldehydes, followed by action of aldehyde dehydrogenase and phosphotransacylase to generate an acyl-phosphate leading to ATP generation in the oxidative branch, while in the reductive branch, an alcohol dehydrogenase is used for cofactor regeneration (Figure 1A and Supplementary Table 1).

**Figure 1.**
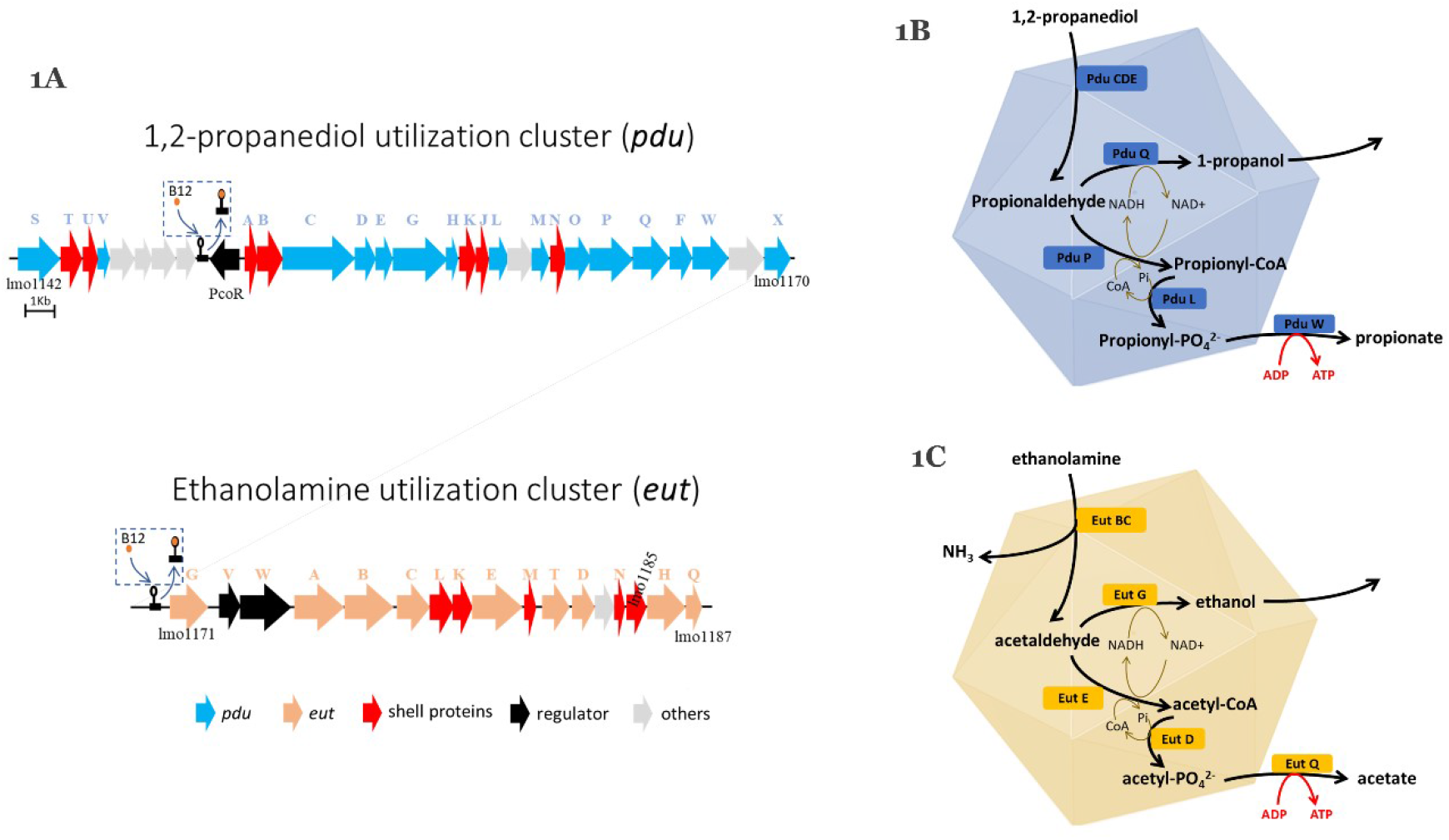
Gene loci organization and core metabolic reactions of bacterial microcompartments (BMCs) in *Listeria monocytogenes* EGDe. **(A)** Gene loci organization of 1,2-propanediol utilization cluster (*pdu*) and ethanolamine utilization cluster (*eut*). Letters above the gene characters indicate the gene name for *pdu* and *eut* separately, see text for details. Vitamin B12-dependent riboswitches to activate *pdu* and *eut* are shown in box. Genes name and proteins ID are in Supplementary Table1. **(B)** *pdu* BMC for 1,2-propanediol utilization. PduCDE, diol dehydratase; PduQ, propanol dehydrogenase. PduP, CoA-dependent propionaldehyde dehydrogenase; PduL, Phosphotransacylase; PduW, propionate kinase. **(C)** *eut* BMC for ethanolamine utilization. EutBC, ethanolamine ammonia lyase; EutG, alcohol dehydrogenase; EutE, acetaldehyde dehydrogenase; EutD, phosphotransacetylase; EutQ, acetate kinase.

### 6.3.2 Activation of *L. monocytogenes* BMC-dependent *pdu* and *eut*

Growth performance, metabolism, transmission electron microscopy and proteomics confirmed B12-dependent BMC activation in *L. monocytogenes pdu* and *eut*-induced cells (data not shown), in line with previous observations [23, 24]. Comparative proteomic profiling of *pdu* and *eut* BMC induced and respective non-induced cells enabled identification of differentially expressed proteins linked to metabolism, stress response and virulence. Analysis of the complete list of identified proteins and subsequent Student’s t-test difference scores of *pdu* induced compared to *pdu* non-induced control cells (Supplementary Table 2), resulted in a selection of 1444 proteins shown in a volcano plot (Figure 2A), with 160 proteins upregulated more than two fold in *pdu* induced cells including 20 proteins encoded in the Pdu cluster, Pdu STUVABCDEGHKJMNOPQFX (Figure 2A blue dots, details in Supplementary Table 2). Comparing *eut* induced cells with *eut* non-induced cells, 161 proteins were upregulated more than 2 fold in *eut* induced cells with in total 1891 identified proteins (Figure 2B blue dots, details in Supplementary Table 3). All of 13 Eut proteins in the *eut* cluster were significant upregulated in *eut* induced cells compared to *eut* non-induced cells. The increased expression of the pdu and eut-encoded proteins in the tested conditions is in line with previously published results [23, 24]. Analysis of the differentially expressed proteins in *pdu* induced and *eut* induced cells shows shared and specific responses that will be discussed in the next section.

**Figure 2.**
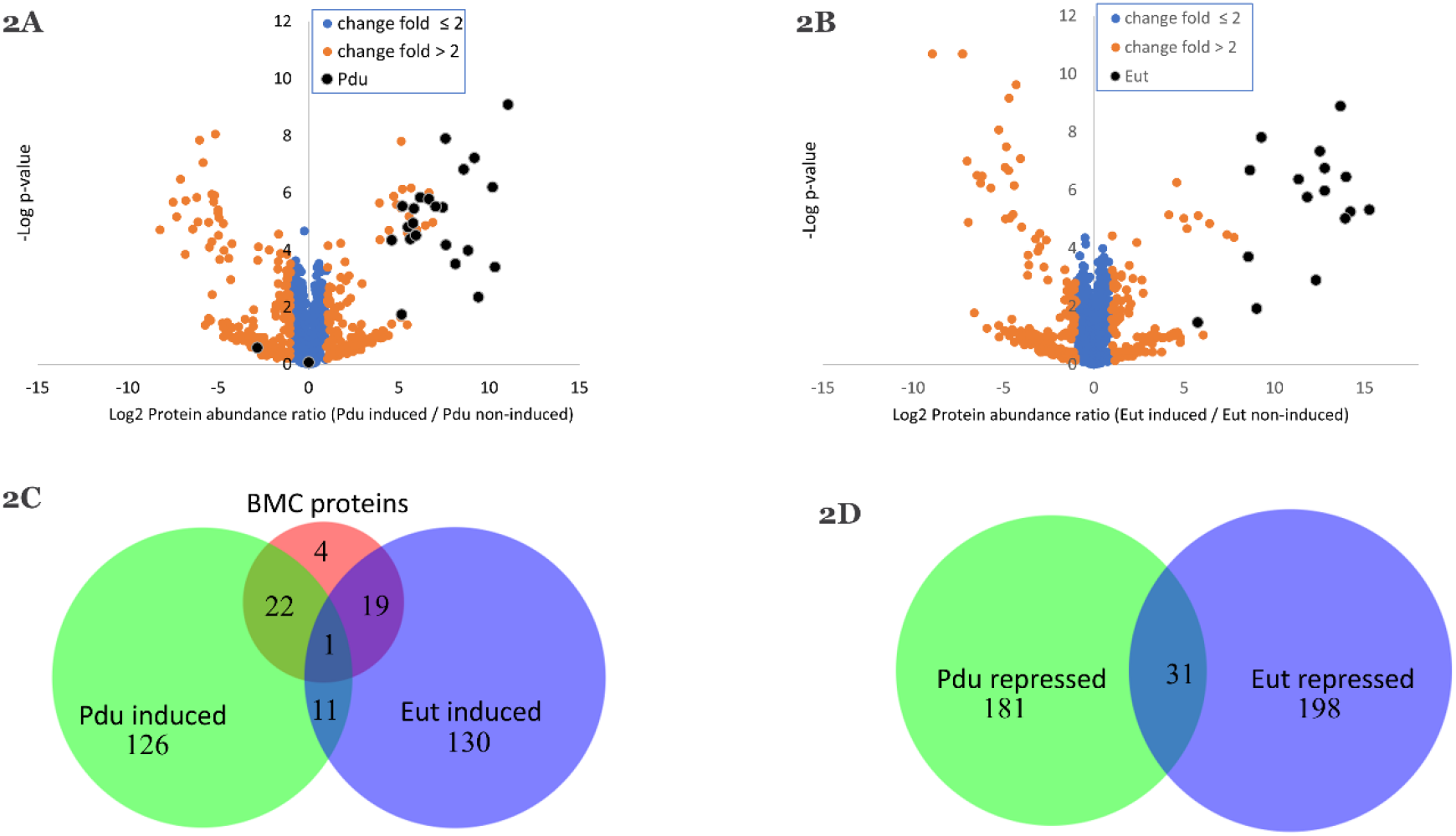
Proteomics analysis of BMC induced and BMC non-induced *Listeria monocytogenes* EGDe. **(A)** Proteomic volcano plot of *pdu* induced cells (Pd + B12 added) compared to *pdu* noninduced cells (Pd added only), full list of proteins is in Supplementary Table 2. **(B)** Proteomic volcano plot of *eut* induced cells (EA + B12 added) compared to *eut* noninduced cells (EA added only), full list of proteins is in Supplementary Table 3, this dataset has been retrieved from Zeng Z, et al, 2021 [24]. In (A) and (B) proteins with the fold change ≤ 2 are in blue, proteins with the fold change > 2 are in orange; Pdu or Eut proteins are indicated in dark blue. **(C)** Venn diagram of the overlapping *pdu* and *eut* induced proteins and **(D)** Venn diagram of the overlapping *pdu* and *eut* repressed proteins. Details of the proteins in (C) and (D) are in Supplementary Table 4 and Supplementary Table 5.

### 6.3.3 Comparative proteome analysis of *pdu* and *eut* BMC induced cells shows overlap in induced and repressed proteins

Shared and specific responses of *pdu* and *eut* BMC induced cells and non-induced cells are shown in Figure 2C and 2D. Shared induced proteins are shown in Figure 2C, with 160 proteins upregulated more than two fold in *pdu* induced cells and 161 proteins upregulated more than two fold in *eut* induced cells, showing 12 common induced proteins. These 12 proteins include several metabolic enzymes, Acetate kinase 2 (Uniprot ID: Q8Y7V1, Gene ID: *Imo1168*), pyruvate - phosphate di-kinase (Q8Y633, *Imo1867*), putative pyruvate-phosphate di-kinase regulatory protein 2 (Q8Y634, *lmo1866*), putative transaldolase 2 (P66957, *lmo0343*), proteins Q8YA21 (*lmo0345*), Q8YA22 (*lmo0344*) and Q8YA23 (*lmo0343*), together function as transketolase (Supplementary Table 4 and Supplementary Figure 2). Additional shared induced proteins include *lmo2158* encoded protein CsbD (Q929L4), a SigB regulon member involved in osmotic and heat stress defense [32], *lmo0818* encoded protein Q8Y8S6, a putative Ca^2+^ ATPase contributing to metal homeostasis [33, 34], and protein *lmo2323* encoded Gp43 (Q8Y4V7), a protein encoded in the bacteriophage A118 gene cluster [35]. In Figure 2D, 212 proteins downregulated more than two fold in *pdu* induced cells and 229 proteins downregulated more than two fold in *eut* induced cells, show 31 common repressed proteins including protein P0DJM0 Internalin A (*lmo0433*) and protein P34024 phospholipase A (*lmo0201*). The remaining set of proteins includes protein Q8Y7U1, zinc uptake regulator (ZurR), that is predicted to coordinate uptake of zinc from the external environment, and a variety of proteins with diverse putative functions and hypotheticals [36] (Supplementary Table 5 and Supplementary Figure 3). Shared and specific *pdu* and *eut* BMC induced and repressed proteins point to metabolic shifts, stress response and virulence, with selected aspects discussed in more detail in sections below.

### 6.3.4 Activation of *pdu* and *eut* BMC metabolism represses glycolysis in *L. monocytogenes*

To visualize the impact of BMC-associated shifts in metabolism, the identified proteins and expression levels (Figure 3A) are mapped to the glycolysis pathway in *L. monocytogenes* EGDe (Supplementary Table 2). Obviously, enzymes involved in the conversion of glyceraldehyde 3-phosphate to pyruvate (Figure 3A, pointed with red arrow) were mostly repressed in *pdu* induced cells compared with *pdu* non-induced cells. Activation of propanoate metabolism (Supplementary Figure 1) is in accordance with increased expression of *pdu* cluster and degradation of 1,2-propanediol to 1-propanol and propionate. Similar to *pdu* induced cells, *eut* induced cells also show a clear restraint of glycolysis (Figure 3B). The enzymes mediating conversion of glyceraldehyde 3-phosphate to pyruvate (Figure 3B, pointed with red arrow) are downregulated in *eut* induced cells compared with *eut* non-induced cells (Figure 3B). Activation of ethanolamine ammonia lyase EutBC in *eut* induced cells degrades ethanolamine into acetaldehyde inside BMCs, that is metabolized into end products acetate and ethanol.

**Figure 3.**
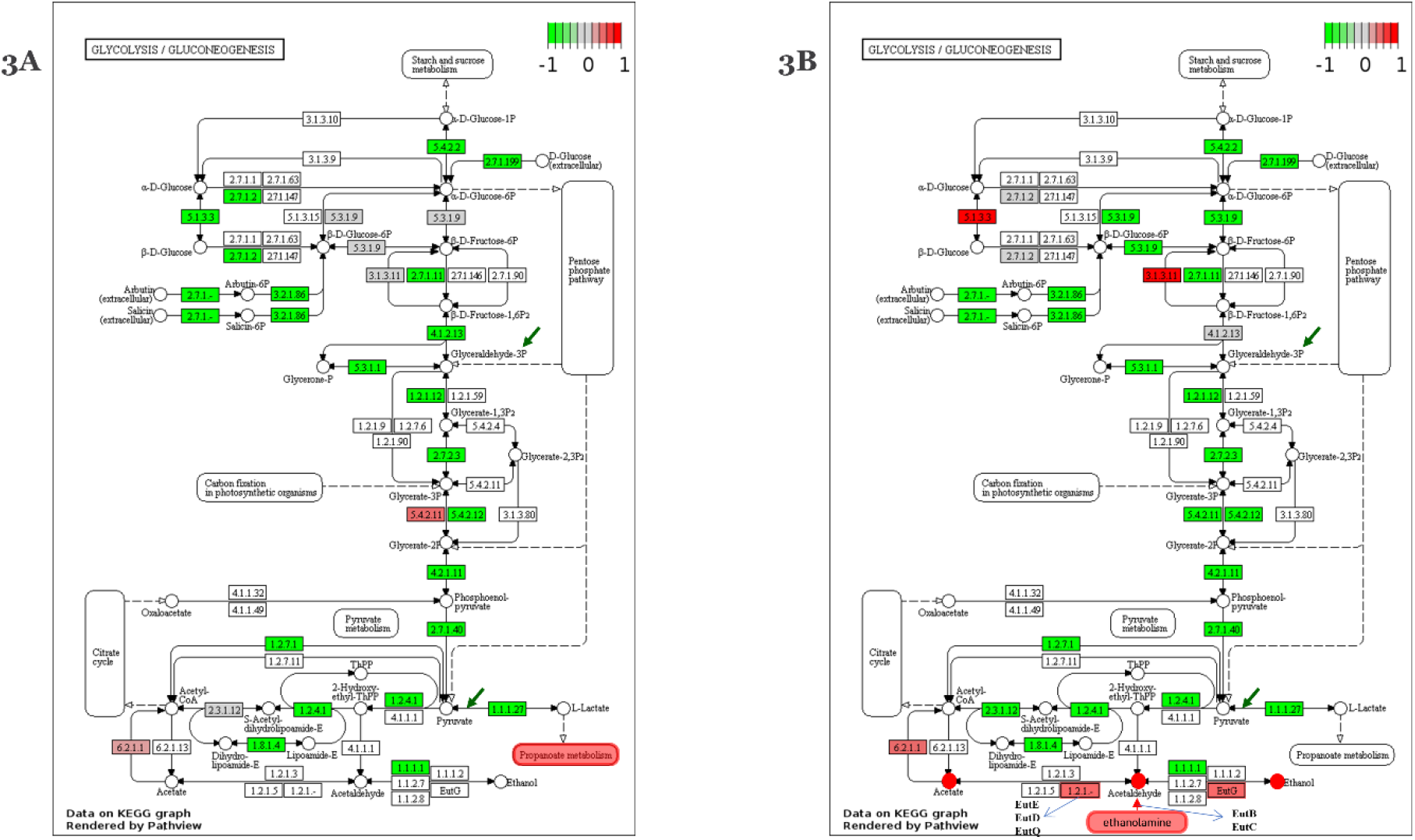
Glycolysis metabolism visualized with proteomic profiling in BMC induced compared BMC non-induced *Listeria monocytogenes* EGDe. **(A)** Glycolysis metabolism visualized with proteomic profiling in *pdu* BMC induced compared with *pdu* BMC non-induced *Listeria monocytogenes* EGDe via Pathview. Details of upregulated propanoate metabolism are shown in Supplementary Figure 1. **(B)** Glycolysis metabolism visualized with proteomic profiling in *eut* BMC induced compared with *eut* BMC non-induced *Listeria monocytogenes* EGDe via Pathview. Upregulated ethanolamine metabolism is customized to be included. In (A) and (B), Colors represent the change intensity of protein expression in BMC induced compared to BMC non-induced *Listeria monocytogenes* EGDe. Lines with arrow represent the metabolic reactions, circles represent metabolites while rectangles represent enzymes with EC number or gene name.

### 6.3.5 *pdu* and *eut* BMC induced stress and virulence associated proteins

Next to shared induced and repressed responses, *pdu* and *eut* specific responses were observed (Supplementary Table 4 for shared induced proteins of *pdu* and *eut* BMC cells, Supplementary Table 5 for shared repressed proteins of *pdu* and *eut* BMC cells). The *pdu* induced cells contained significant higher levels of CtsR, transcript regulator of heat shock stress proteins [37]; Stressosome component Blue-light photoreceptor lmo0799, involved in light and redox control of SigB expression [38]; Sigma B anti-anti-sigma factor, Serine-protein kinase RsbW, GbuB; Sigma B-dependent osmoprotectant betaine transporter [39]; Heat-inducible transcription repressor HrcA [40]; lmo1021, encoding histidine kinase of the LiaSR two-component system (lmo1021 and lmo1022) that plays a role in resistance to antimicrobials [41]; Protease HtpX homolog, involved in membrane/cell envelope stress response [42]; LexA repressor, involved in DNA damage repair, and DPS, DNA protection during starvation protein [42]. In addition, a number of bacteriophage A118 encoded proteins are higher expressed in pdu induced cells, conceivably due to activation of gene expression following DNA damage [35]. No significant higher expression of PrfA or sigmaB controlled virulence factors is observed in *pdu* induced cells.

However, both *pdu* induced and non-induced control cells contain internalin B, Q8Y7G3 encoded putative membrane bound zinc metalloprotease (Lmo1318) involved in release of Q8Y436 peptide pheromone encoding lipoprotein A, PplA (Lmo2637), that conceivably supports activation of PrfA [43, 44], and Listeria Adhesion Protein (LAP, lmo1634), involved in host cell invasion [45], with the latter protein significantly higher expressed in pdu non-induced cells.

The *eut* induced cells express higher levels of Phosphate import ATP-binding proteins PstB 1 and PstB 2 [3]; Phosphate-specific transport system accessory protein PhoU (*lmo2494*) [3]; Manganese transport system membrane protein MntC [3]; OpuCD protein, involved in sigB dependent uptake of osmoprotectant carnitine [46]; Fosfomycin resistance protein FosX, involved in resistance to antimicrobials [47]; ATP-dependent protease ClpE, involved in protein quality control [48]; and RecS protein, ATP-dependent DNA helicase [3]; RecG, and Endonuclease III, all three involved in DNA damage repair [3]. In addition, a number of bacteriophage A118 encoded proteins are higher expressed in *eut* induced cells, conceivably due to activation of gene expression following DNA damage. Concerning activation of virulence factors, the *eut* induced cells showed significantly higher levels of Listeriolysin regulatory protein PrfA and P13128 encoded Listeriolysin O (*hly*). The *eut* induced and control cells also express Q8Y7G3 encoded zinc metalloprotease *(lmo1318)* and Q8Y436 encoded PplA (*lmo2637*), and LAP *(lmo1634)* as described above, and in addition produce P33379 encoded Actin assembly-inducing protein ActA, P34024 encoded 1-phosphatidylinositol phosphodiesterase PlcA; and P33378 encoded Phospholipase B, PlcB, although the latter is significantly higher expressed in *eut* non-induced cells.

These results point to shared stress defense responses in *pdu* and *eut* induced cells including uptake of osmoprotectants (Gbu and OpuC transport proteins), activation of chaperons for quality control (ClpP and ClpE), and enzymes involved in DNA damage repair (LexA and RecS, RecG, recombinase), with a more extensive response in eut cells at the level of regulators and proteins involved in transport of phosphate and transition metals such as manganese (Mn), calcium (Ca), Iron/heme, and cadmium (Cd). Notably, only *eut* induced cells showed significant higher expression of virulence factors PrfA and listeriolysin O (LLO, Hly). Subsequent vitro virulence assays with *pdu* and *eut* BMC primed cells are discussed in the next section.

### 6.3.6 Impact of pdu and eut BMC activation on Caco-2 cell adhesion, invasion and translocation

In vitro adhesion, invasion and translocation assays with Caco-2 cells were performed using primed pdu and eut BMC cells and non-induced control cells Figure 4). As shown in Figure 4A, the ability to translocate Caco-2 cells monolayers in a trans-well system, is significantly higher for *pdu* induced *L. monocytogenes* cells compared to *pdu* non-induced cells (0.5 log of CFU/well improvement), while no significant differences were observed for adhesion and invasion efficacy (Figure 4A). Similar to *pdu* induced cells, *eut* induced cells show significantly higher translocation efficacy (0.6 log of CFU/well increase), while no significant improvement of adhesion and invasion capacity is observed (Figure 4B). Our results provide evidence for enhanced Caco-2 cell monolayer translocation efficacy of pdu and eut BMC primed cells compared to non-induced control cells.

**Figure 4.**
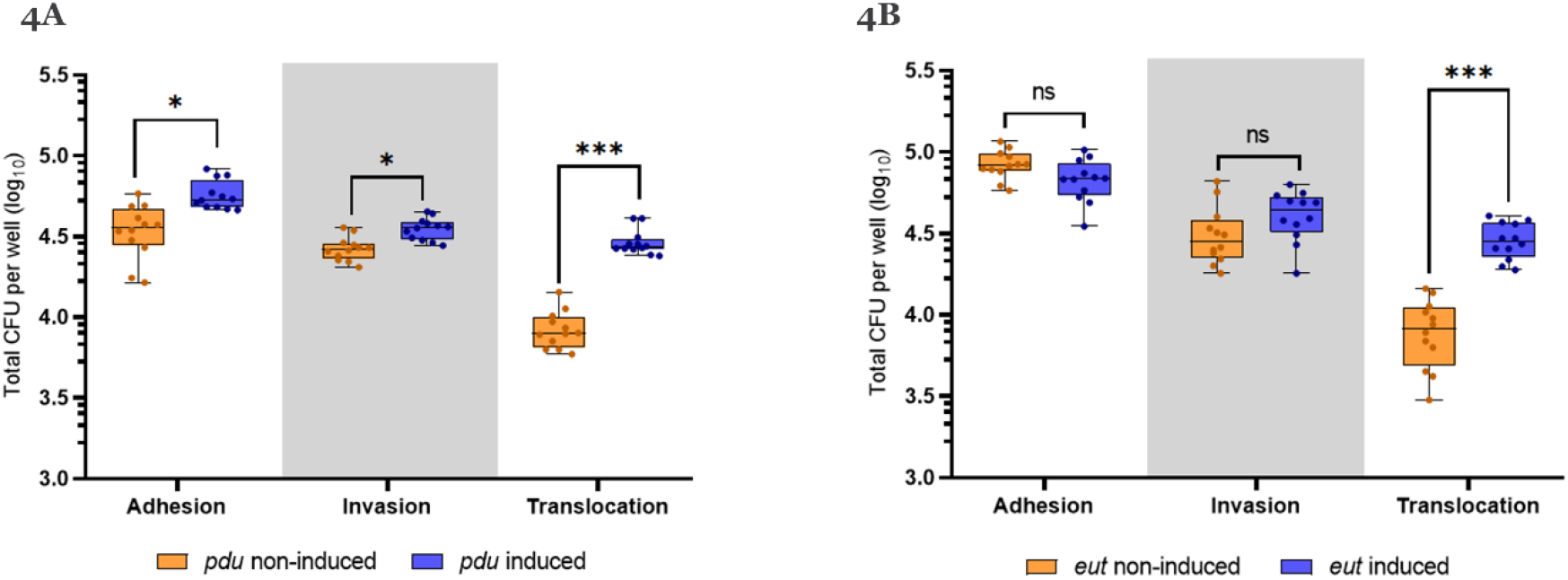
Caco-2 cell assays with BMC induced and BMC non-induced *Listeria monocytogenes* EGDe. **(A)** is for *pdu* BMC induced and *pdu* BMC non-induced cells. **(B)** is for *eut* BMC induced and *eut* BMC non-induced cells. X axis show three groups of columns which are adhesion assay, invasion assay and translocation assay. Y axis is the total CFU per well in log_10_ after the assays while initial inputs for adhesion and invasion are approximate 6.8 log_10_ CFU/well and the initial inputs for translocation are approximate 6.5 log_10_ CFU/well. Statistical significance are indicated (*** means P≤0.001; * means P≤0.05; ns means > 0.05 by Holm-Sidak T-test)

## 6.4 Discussion

Bacterial microcompartments (BMCs) are proteinaceous organelles that optimize specific metabolic pathways referred to as metabolosomes involving transient production of toxic volatile metabolites such as aldehydes [17, 49]. We recently provided evidence for a role of BMC-dependent utilization of propanediol *(pdu)* and ethanolamine (*eut*) in *L. monocytogenes* resulting in growth stimulation under anaerobic conditions [23, 24]. In this study we performed a comparative proteomics analysis of *pdu* and *eut* BMC primed L. *monocytogenes* cells and non-induced control cells, to identify specific and shared responses focusing on metabolic, stress response and virulence parameters, and to correlate these to adhesion, invasion and translocation efficacy in caco-2 assays. Expression analysis of the complete lists of identified proteins showed 160 of 1444 proteins upregulated more than two fold in *pdu* induced cells including 20 proteins encoded in the *pdu* cluster, and 161 of 1891 identified proteins in *eut* induced cells, including all 13 proteins encoded in the eut cluster. Creation of BMCs with key metabolic turnover steps of *pdu* and *eut* enzymes encapsulated, occurs in a stepwise manner [13, 14], thereby creating a protection against the respective toxic intermediates propionaldehyde and acetaldehyde. Proteomics analysis of shared and specific *pdu* BMC and *eut* BMC responses indicates metabolic shifts and activation of stress defense proteins, conceivably as a response to aldehyde-induced redox stress before BMC production is completed. Shared main themes in *pdu* and *eut* BMC primed cells include expression of transcriptional regulators and/or respective regulon members of ZurR, a transcriptional repressor of zinc (Zn) transporters [50]; SigB, CtsR, and HrcA, transcriptional activators/repressors involved in coordinating cellular stress defense including uptake systems for osmoprotectants and synthesis of a range of chaperons contributing to maintenance of protein quality and functionality [51], and RecA/LexA, regulators of *L. monocytogenes* SOS response including enzymes involved in DNA damage repair [52, 53]. The latter stress phenomenon is conceivable linked to expression activation of *L. monocytogenes* EGDe prophage A118 [35], with highest relative abundance of A118 GP43 protein (Lmo2323) in both *pdu* and *eut* primed cells. Expression of additional phage proteins was noted, but holin and lysin proteins were not detected (Supplementary Table 1), in line with the observation that *pdu* and *eut* induced cells reach high cell counts during anaerobic growth indicating no or very limited lysis of cells (data not show, [23, 24]). Notably, previous studies showed that the A118 gene cluster is inserted in the *L. monocytogenes* EGDe *comK* gene, which renders it inactive in cells grown at 30°C, while excision of the cluster leading to expression of the *L. monocytogenes comK* gene and corresponding competence genes at 37°C, contributes to intracellular growth following bacterial escape from macrophage phagosomes [54, 55]. Notably, activation of SOS response is another facet of *pdu* and *eut* BMC priming that can affect efficacy and timing of the BMC encased pathways with concomitant aldehyde production, an aspect that has been largely ignored up to now including possible activation of lytic or lysogenic phages, that may affect competitive fitness of producer cells.

Analysis of specific responses shows a higher expression in *eut* induced cells of proteins involved in transport of phosphate and transition metals such as manganese (Mn), calcium (Ca), Iron/heme, and cadmium (Cd). This might point to increased acetaldehyde-induced oxidative stress affecting transition metal homeostasis leading to activation of DNA-binding metal-responsive transcription factors, metal-acquisition or secretion, metal-detoxification and metal-storage systems, some of which represent key *L. monocytogenes* virulence determinants [56]. In addition, BMC-dependent metabolism may be affected by availability of compounds including Fe^3+^ that can act as extracellular electron acceptors allowing a shift towards the ATP generating oxidative branch resulting in higher cell numbers, as we previously showed for eut BMC primed *L. monocytogenes* cells anaerobically grown in defined medium without and with added Fe^3+^ [24]. A physiological link between the large number of differentially expressed metal transport systems and significant higher expression of PrfA and listeriolysin (LLO) in BMC *eut* induced cells, may be explained by recent findings reported by Gaballa et al. (2021) [57], about the role of divalent metal ion scavengers Chelex and activated charcoal in the induction of PrfA regulon expression in complex medium. This study showed that the expression of the PrfA-regulated gene *hly*, which encodes listeriolysin O (LLO), is induced 5- and 8-fold in *L. monocytogenes* cells grown in Chelex-treated BHI (Ch-BHI) and in the presence of activated charcoal (AC-BHI), respectively, relative to cells grown in BHI medium. Additional experiments provide evidence that intracellular metal depletion contributes to activation of PrfA and that genes in its regulon are differentially expressed with LLO showing highest modulation. The authors suggest that metal ion abundance plays a role in modulating expression of PrfA-regulated virulence genes in *L. monocytogenes* [49]. Additional experiments are required to confirm the hypothesis that increased expression of metal transporters in *eut* BMC induced cells signifies a homeostatic response to metal depletion, concomitant with increased expression of PrfA and LLO in eut BMC induced cells.

Comparative analysis of Caco-2 cell monolayer adhesion, invasion and translocation, showed similar adhesion and invasion efficacy of *pdu* and *eut* non-induced compared to *pdu* and *eut* BMC primed cells, with the latter BMC activated cells showing significantly higher translocation efficacy. Comparative analysis of virulence factor expression, shows specific activation of PrfA and listeriolysin LLO in *eut* induced cells but not in *pdu* induced cells, while no increased expression of *L. monocytogenes* virulence factors Internalin A (InlA), Internalin B (InlB), Phospholipase A and B (PlcA, PlcB), and Actin (ActA), is observed. Analysis of the complete list of proteins in *eut* non-induced and *eut* induced cells (Supplementary Table 3), showed the presence of Actin assembly-inducing protein ActA, 1-phosphatidylinositol phosphodiesterase PlcA, and Phospholipase B, PlcB, in both types of cells. It is therefore possible that enhanced translocation of *eut* induced cells is not solely linked to activation of PrfA and more importantly to LLO activity, but is supported by activity of PlcA, PlcB and ActA. To confirm roles of LLO, that has been shown to mediate alternative routes of invasion and translocation and possible contributions by other virulence factors in enhanced translocation efficacy of *eut* induced cells [12, 43, 44, 58], further studies are required. Obviously, the same holds for identification of factors that contribute to enhanced Caco-2 cell monolayer translocation of pdu induced cells. Notably, analysis of the complete list of proteins present in *pdu* non-induced and induced cells (Supplementary Table 2), shows the presence of internalin B, next to putative membrane bound zinc metalloprotease *(lmo1318)* involved in release of peptide pheromone encoding lipoprotein A, PplA (*lmo2637*), that conceivably supports activation of PrfA [43, 57], and Aldehyde-alcohol dehydrogenase/Listeria Adhesion Protein (LAP, *lmo1634*) . Notably, with the exception of InlB, these proteins are also present in eut non induced and induced cells, with LAP, previously shown to provide an alternative route of entry and translocation [10, 11]. Whether these virulence factors in combination with other cellular and physiological features of *pdu* BMC primed cells contribute to enhanced translocation efficacy remains to be confirmed.

In conclusion, we have provided evidence for shared and specific metabolic, stress resistance and virulence associated responses at proteome level in pdu and eut BMC primed *L. monocytogenes* cells. Shared stress responses include activation of stress defense proteins and SOS DNA damage repair enzymes [52, 53], conceivably linked to activation and selected expression of previously described lysogenic prophage A118 gene cluster. Most *L. monocytogenes* strains carry prophages in their genome, including A118 and A118-like prophages such as in *L. monocytogenes* 10403S [35, 54, 55]. Using *L. monocytogenes* 10403S as a model, Pasechnek et al. [55] recently provided evidence for temperature-dependent activation of lysogeny as a bacteria-phage adaptive response in the mammalian environment, with temperature as a crucial factor controlling excision of the phage gene cluster and expression of selected phage proteins that excluded the holin and lysin proteins, associated with bacterial host cell lysis. Since in our studies we used *L. monocytogenes* EGDe grown at 30°C, mimicking priming of *pdu* and *eut* BMCs before entering the human host for example during transmission in foods, future experiments should also include cells primed at 37°C to mimic activation of BMCs inside the mammalian host. Inclusion of multiple *L. monocytogenes* strains, without and with lysogenic prophages, will further add to our understanding of mechanisms underlying production and functionality of pdu and eut BMCs in *L. monocytogenes* competitive fitness and survival during transmission in the food chain and inside the mammalian host.

## 6.5 Supplementary Materials

**Supplementary figure 1.**
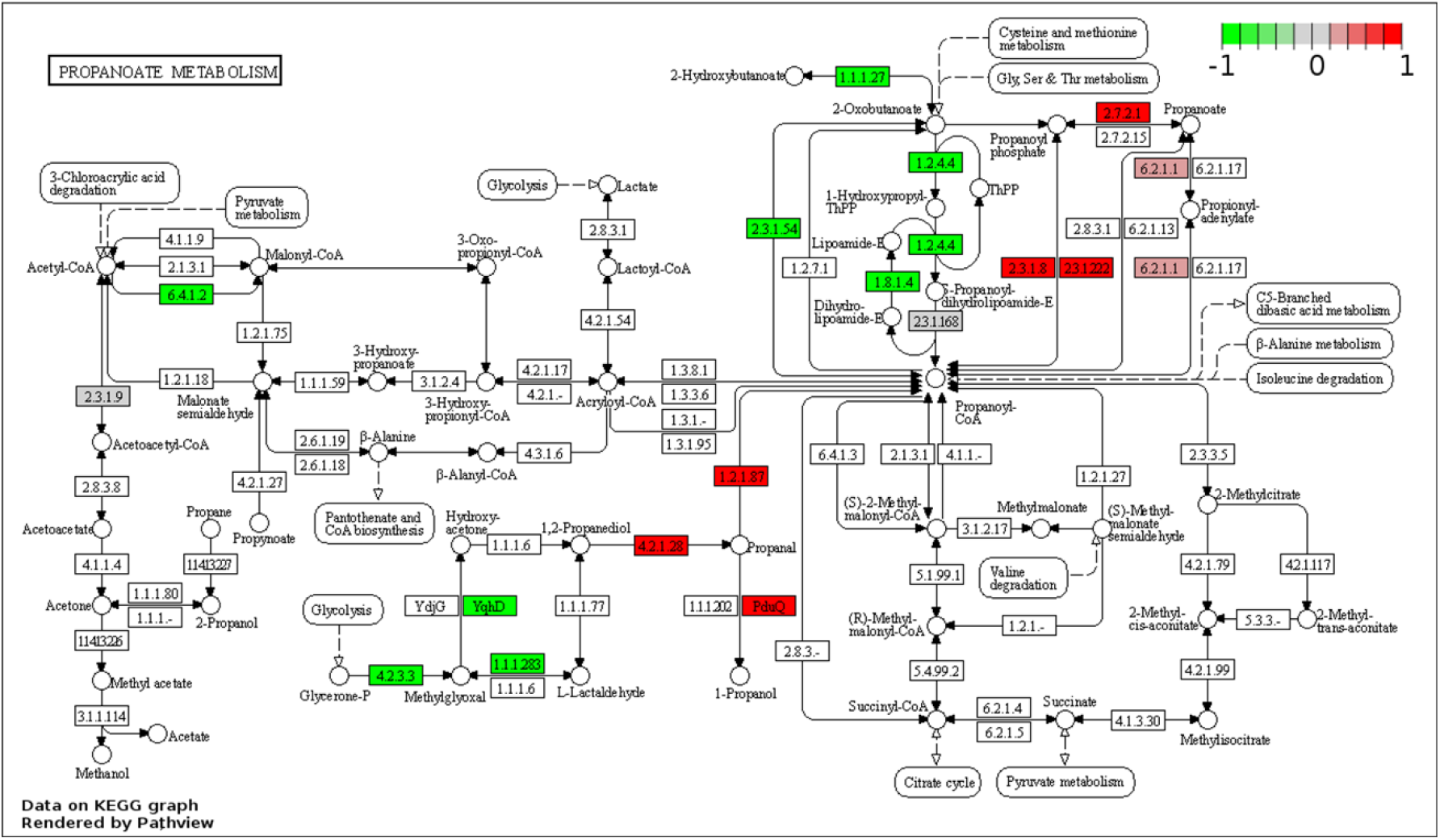
Propanoate metabolism visualized with proteomic profiling in *pdu* BMC induced compared *pdu* BMC non-induced *Listeria monocytogenes* EGDe. Colors represent the change intensity of protein expression in *pdu* BMC induced compared to *pdu* BMC non-induced *Listeria monocytogenes* EGDe. Lines with arrow represent the metabolic reactions, circles represent metabolites while rectangles represent enzymes with EC number or gene name.

**Supplementary figure 2.**
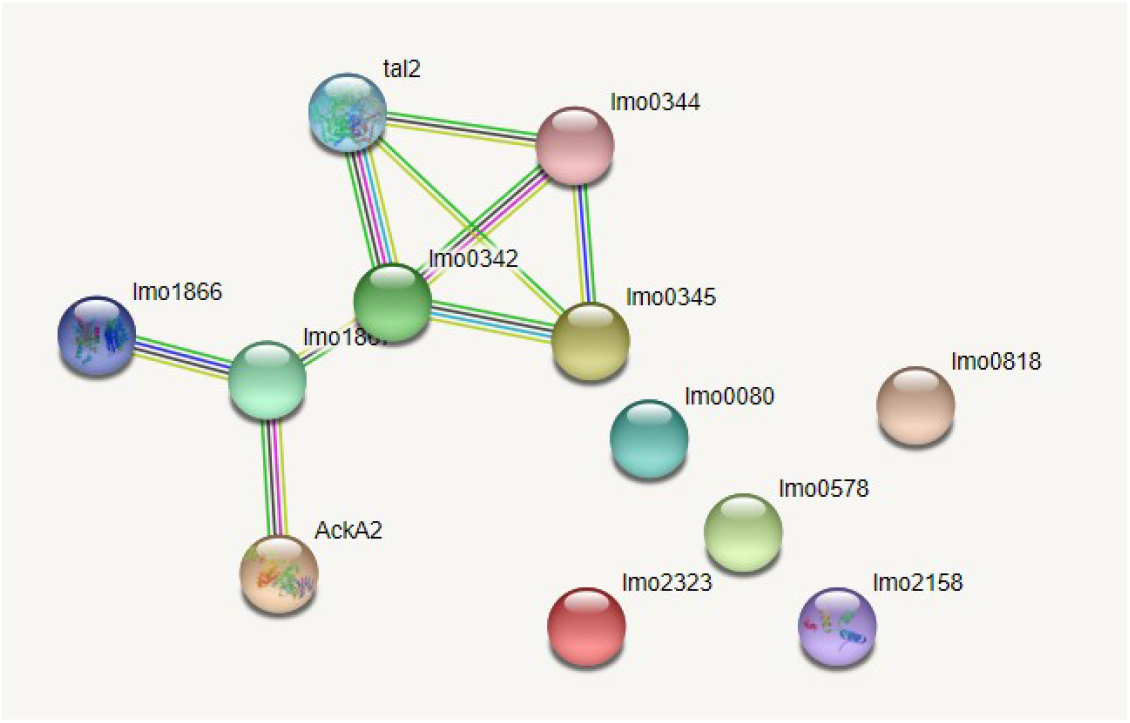
STRING protein-protein interactions of common induced 12 proteins in pdu-induced cells and eut-induced cells.

**Supplementary figure 3.**
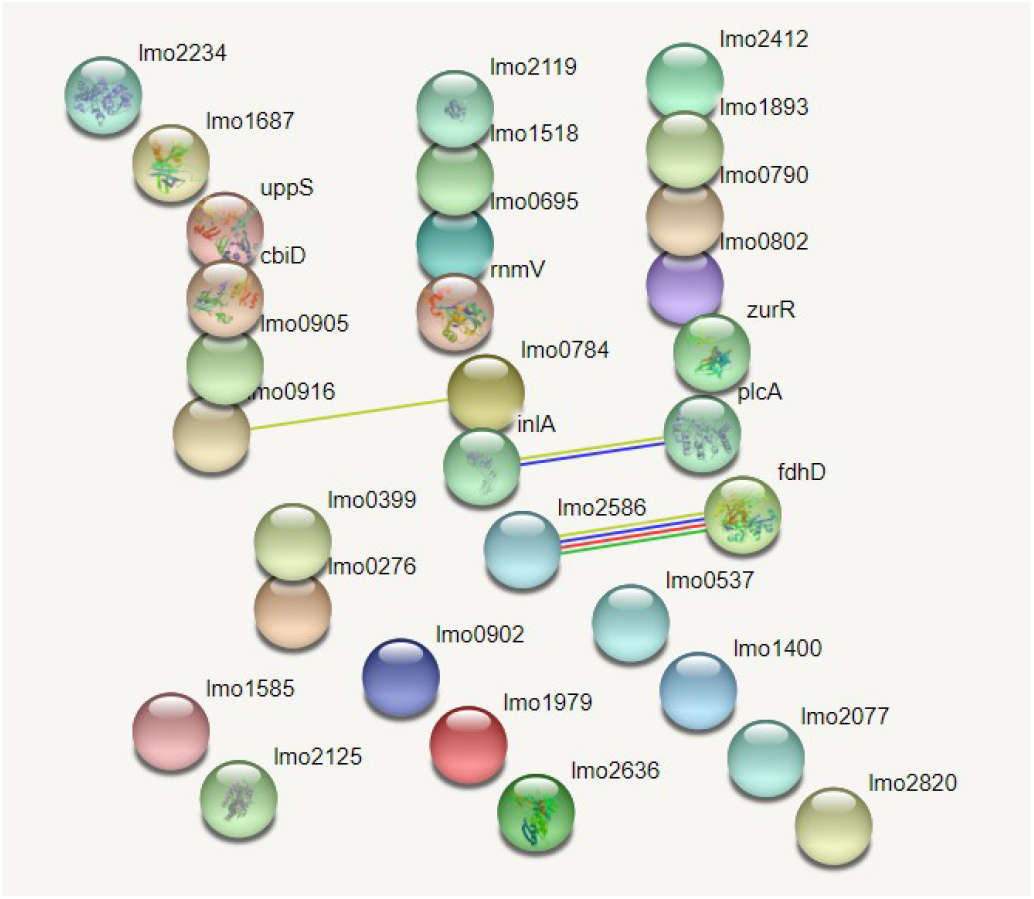
STRING protein-protein interactions of common repressed 31 proteins in pdu-induced cells and eut-induced cells.

**Supplementary Table 1**. Gene annotation, proteins IDs and protein domain anotation of *pdu* and *eut* cluster

**Supplementary Table 2**. Protein profiling of *pdu* induced cells compared to *pdu* non-induced cells

**Supplementary Table 3**. Protein profiling of *eut* induced cells compared to *eut* non-induced cells

**Supplementary Table 4**. Shared induced proteins of *pdu* and *eut* BMC cells.

**Supplementary Table 5**. Shared repressed proteins of *pdu* and *eut* BMC cells.

The table lists UniProt protein ID, X value representing Log2 Protein abundance ratio (eut induced/non-induced in LB), Y value representing -Log10 p-value (induced condition /non-induced condition), NCBI protein Annotation and NCBI Gene ID.

## References

1. Radoshevich, L. and P. Cossart, Listeria monocytogenes: towards a complete picture of its physiology and pathogenesis. Nature Reviews Microbiology, 2018. 16(1): p. 32.

2. Gandhi, M. and M.L. Chikindas, Listeria: a foodborne pathogen that knows how to survive. International journal of food microbiology, 2007. 113(1): p. 1–15.

3. Freitag, N.E., G.C. Port, and M.D. Miner, Listeria monocytogenes—from saprophyte to intracellular pathogen. Nature Reviews Microbiology, 2009. 7(9): p. 623–628.

4. Desai, A.N., et al., Changing epidemiology of Listeria monocytogenes outbreaks, sporadic cases, and recalls globally: A review of ProMED reports from 1996 to 2018. International Journal of Infectious Diseases, 2019. 84: p. 48–53.

5. Tasara, T. and R. Stephan, Cold stress tolerance of Listeria monocytogenes: a review of molecular adaptive mechanisms and food safety implications. Journal of food protection, 2006. 69(6): p. 1473–1484.

6. Portnoy, D.A., V. Auerbuch, and I.J. Glomski, The cell biology of Listeria monocytogenes infection: the intersection of bacterial pathogenesis and cell-mediated immunity. The Journal of cell biology, 2002. 158(3): p. 409–414.

7. Ragon, M., et al., A new perspective on Listeria monocytogenes evolution. PLoS Pathogens, 2008. 4(9): p. e1000146.

8. Mengaud, J., et al., E-cadherin is the receptor for internalin, a surface protein required for entry of L. monocytogenes into epithelial cells. Cell, 1996. 84(6): p. 923–932.

9. Tattoli, I., et al., Listeria phospholipases subvert host autophagic defenses by stalling pre-autophagosomal structures. The EMBO journal, 2013. 32(23): p. 3066–3078.

10. Horn, N. and A.K. Bhunia, Food-associated stress primes foodborne pathogens for the gastrointestinal phase of infection. Frontiers in microbiology, 2018. 9: p. 1962.

11. Kim, H. and A.K. Bhunia, Secreted Listeria adhesion protein (Lap) influences Lap-mediated Listeria monocytogenes paracellular translocation through epithelial barrier. Gut pathogens, 2013. 5(1): p. 1–11.

12. Nguyen, B.N., B.N. Peterson, and D.A. Portnoy, Listeriolysin O: a phagosome-specific cytolysin revisited. Cellular microbiology, 2019. 21(3): p. e12988.

13. Jakobson, C.M. and D. Tullman-Ercek, Dumpster diving in the gut: bacterial microcompartments as part of a host-associated lifestyle. PLoS Pathogens, 2016. 12(5): p. e1005558.

14. Kerfeld, C.A., et al., Bacterial microcompartments. Nature Reviews Microbiology, 2018. 6(5): p. 227.

15. Thiennimitr, P., et al., Intestinal inflammation allows Salmonella to use ethanolamine to compete with the microbiota. Proceedings of the National Academy of Sciences, 2011. 108(42): p. 17480–17485.

16. Kerfeld, C.A. and O. Erbilgin, Bacterial microcompartments and the modular construction of microbial metabolism. Trends in microbiology, 2015. 23(1): p. 22–34.

17. Prentice, M.B., Bacterial microcompartments and their role in pathogenicity. Current Opinion in Microbiology, 2021. 63: p. 19–28.

18. Agus, A., K. Clément, and H. Sokol, Gut microbiota-derived metabolites as central regulators in metabolic disorders. Gut, 2021. 70(6): p. 1174–1182.

19. Anast, J.M., T.A. Bobik, and S. Schmitz-Esser, The Cobalamin-dependent gene cluster of Listeria monocytogenes: implications for virulence, stress response, and food safety. Frontiers in microbiology, 2020. 11.

20. Rowley, C.A., C.J. Anderson, and M.M. Kendall, Ethanolamine influences human commensal Escherichia coli growth, gene expression, and competition with enterohemorrhagic E. coli O157: H7. mBio, 2018. 9(5): p. e01429–18.

21. Tang, S., et al., Transcriptomic analysis of the adaptation of Listeria monocytogenes to growth on vacuum-packed cold smoked salmon. Appl. Environ. Microbiol., 2015. 81(19): p. 6812–6824.

22. Schardt, J., et al., Comparison between Listeria sensu stricto and Listeria sensu lato strains identifies novel determinants involved in infection. Scientific reports, 2017. 7(1): p. 17821.

23. Zeng, Z., et al., Bacterial microcompartment-dependent 1, 2-propanediol utilization stimulates anaerobic growth of Listeria monocytogenes EGDe. Frontiers in Microbiology, 2019. 10: p. 2660.

24. Zeng, Z., et al., Bacterial Microcompartments Coupled with Extracellular Electron Transfer Drive the Anaerobic Utilization of Ethanolamine in Listeria monocytogenes. mSystems, 2021. 6(2).

25. Mellin, J., et al., A riboswitch-regulated antisense RNA in Listeria monocytogenes. Proceedings of the National Academy of Sciences, 2013. 110(32): p. 13132–13137.

26. Mellin, J., et al., Sequestration of a two-component response regulator by a riboswitch-regulated noncoding RNA. Science, 2014. 345(6199): p. 940–943.

27. Vizcaino, J.A., et al., 2016 update of the PRIDE database and its related tools (vol 44, pg D447, 2016). Nucleic Acids Research, 2016. 44(22): p. 11033–11033.

28. Mistry, J., et al., Challenges in homology search: HMMER3 and convergent evolution of coiled-coil regions. Nucleic acids research, 2013. 41(12): p. e121–e121.

29. Hulsen, T., J. de Vlieg, and W. Alkema, BioVenn-a web application for the comparison and visualization of biological lists using area-proportional Venn diagrams. BMC genomics, 2008. 9(1): p. 1–6.

30. Szklarczyk, D., et al., STRING v11: protein-protein association networks with increased coverage, supporting functional discovery in genome-wide experimental datasets. Nucleic acids research, 2019. 47(D1): p. D607–D613.

31. Luo, W. and C. Brouwer, Pathview: an R/Bioconductor package for pathway-based data integration and visualization. Bioinformatics, 2013. 29(14): p. 1830–1831.

32. Kragh, M.L. and L.T. Hansen, Initial Transcriptomic Response and Adaption of Listeria monocytogenes to Desiccation on Food Grade Stainless Steel. Frontiers in Microbiology, 2020. 10.

33. Morth, J.P. and K.L. Hein, Listeria Monocytogenes Lmo0818-Exploring a Putative Ca2+-ATPase, to Understand Calcium Ion Specificity. Biophysical Journal, 2013. 104(2): p. 300A–300A.

34. Jesse, H.E., I.S. Roberts, and J.S. Cavet, Metal Ion Homeostasis in Listeria monocytogenes and Importance in Host-Pathogen Interactions, in Advances in Microbial Physiology, Vol 65: Advances in Bacterial Pathogen Biology, R.K. Poole, Editor. 2014. p. 83–123.

35. Loessner, M.J., et al., Complete nucleotide sequence, molecular analysis and genome structure of bacteriophage A118 of Listeria monocytogenes: implications for phage evolution. Molecular Microbiology, 2000. 35(2): p. 324–340.

36. Dowd, G.C., et al., Investigation of the role of ZurR in the physiology and pathogenesis of Listeria monocytogenes. Fems Microbiology Letters, 2012. 327(2): p. 118–125.

37. Nair, S., et al., CtsR controls class III heat shock gene expression in the human pathogen Listeria monocytogenes. Molecular Microbiology, 2000. 35(4): p. 800–811.

38. O’Donoghue, B., et al., Blue-Light Inhibition of Listeria monocytogenes Growth Is Mediated by Reactive Oxygen Species and Is Influenced by sigma(B) and the Blue-Light Sensor Lmo0799. Applied and Environmental Microbiology, 2016. 82(13): p. 4017–4027.

39. Guerreiro, D.N., T. Arcari, and C.P. O’Byrne, The sigma(B)-Mediated General Stress Response of Listeria monocytogenes: Life and Death Decision Making in a Pathogen. Frontiers in Microbiology, 2020. 11.

40. Hu, Y., et al., Transcriptomic and phenotypic analyses suggest a network between the transcriptional regulators HrcA and sigma(B) in Listeria monocytogenes. Applied and Environmental Microbiology, 2007. 73(24): p. 7981–7991.

41. Collins, B., et al., Assessing the Contributions of the LiaS Histidine Kinase to the Innate Resistance of Listeria monocytogenes to Nisin, Cephalosporins, and Disinfectants. Applied and Environmental Microbiology, 2012. 78(8): p. 2923–2929.

42. Arolas, J.L., et al., Expression and purification of integral membrane metallopeptidase HtpX. Protein Expression and Purification, 2014. 99: p. 113–118.

43. de las Heras, A., et al., Regulation of Listeria virulence: PrfA master and commander. Current opinion in microbiology, 2011. 14(2): p. 118–127.

44. Scortti, M., et al., The PrfA virulence regulon. Microbes and Infection, 2007. 9(10): p. 1196–1207.

45. dos Santos, P.T., et al., Listeria monocytogenes Relies on the Heme-Regulated Transporter hrtAB to Resist Heme Toxicity and Uses Heme as a Signal to Induce Transcription of Imo1634, Encoding Listeria Adhesion Protein. Frontiers in Microbiology, 2018. 9.

46. Fraser, K.R., et al., Identification and characterization of an ATP binding cassette L-carnitine transporter in Listeria monocytogenes. Applied and Environmental Microbiology, 2000. 66(11): p. 4696–4704.

47. Fillgrove, K.L., et al., Structure and mechanism of the genomically encoded fosfomycin resistance protein, FosX, from Listeria monocytogenes. Biochemistry, 2007. 46(27): p. 8110–8120.

48. Nair, S., E. Milohanic, and P. Berche, ClpC ATPase is required for cell adhesion and invasion of Listeria monocytogenes. Infection and Immunity, 2000. 68(12): p. 7061–7068.

49. Kerfeld, C.A., et al., Bacterial microcompartments. Nature Reviews Microbiology, 2018.

50. Dowd, G.C., et al., Investigation of the role of ZurR in the physiology and pathogenesis of Listeria monocytogenes. FEMS microbiology letters, 2012. 327(2): p. 118–125.

51. Chaturongakul, S., et al., Transcriptomic and Phenotypic Analyses Identify Coregulated, Overlapping Regulons among PrfA, CtsR, HrcA, and the Alternative Sigma Factors sigma(B), sigma(C), sigma(H), and sigma(L) in Listeria monocytogenes. Applied and Environmental Microbiology, 2011. 77(1): p. 187–200.

52. van der Veen, S., et al., The SOS response of Listeria monocytogenes is involved in stress resistance and mutagenesis. Microbiology-Sgm, 2010. 156: p. 374–384.

53. van der Veen, S., et al., The heat-shock response of Listeria monocytogenes comprises genes involved in heat shock, cell division, cell wall synthesis, and the SOS response. Microbiology-Sgm, 2007. 153: p. 3593–3607.

54. Rabinovich, L., et al., Prophage Excision Activates Listeria Competence Genes that Promote Phagosomal Escape and Virulence. Cell, 2012. 150(4): p. 792–802.

55. Pasechnek, A., et al., Active Lysogeny in Listeria Monocytogenes Is a Bacteria-Phage Adaptive Response in the Mammalian Environment. Cell Reports, 2020. 32(4).

56. Jesse, H.E., I.S. Roberts, and J.S. Cavet, Metal ion homeostasis in Listeria monocytogenes and importance in host-pathogen interactions. Advances in microbial physiology, 2014. 65: p. 83–123.

57. Gaballa, A., et al., Characterization of the roles of activated charcoal and Chelex in the induction of PrfA regulon expression in complex medium. Plos one, 2021. 16(4): p. e0250989.

58. Nadon, C.A., et al., Sigma B contributes to PrfA-mediated virulence in Listeria monocytogenes. Infection and immunity, 2002. 70(7): p. 3948–3952.

